# WESTPA 2.0: High-performance upgrades for weighted ensemble simulations and analysis of longer-timescale applications

**DOI:** 10.1101/2021.12.05.471280

**Authors:** John D. Russo, She Zhang, Jeremy M. G. Leung, Anthony T. Bogetti, Jeff P. Thompson, Alex J. DeGrave, Paul A. Torrillo, A. J. Pratt, Kim F. Wong, Junchao Xia, Jeremy Copperman, Joshua L. Adelman, Matthew C. Zwier, David N. LeBard, Daniel M. Zuckerman, Lillian T. Chong

## Abstract

The weighted ensemble (WE) family of methods is one of several statistical-mechanics based path sampling strategies that can provide estimates of key observables (rate constants, pathways) using a fraction of the time required by direct simulation methods such as molecular dynamics or discrete-state stochastic algorithms. WE methods oversee numerous parallel trajectories using intermittent overhead operations at fixed time intervals, enabling facile interoperability with any dynamics engine. Here, we report on major upgrades to the WESTPA software package, an open-source, high-performance framework that implements both basic and recently developed WE methods. These upgrades offer substantial improvements over traditional WE. Key features of the new WESTPA 2.0 software enhance efficiency and ease of use: an adaptive binning scheme for more efficient surmounting of large free energy barriers, streamlined handling of large simulation datasets, exponentially improved analysis of kinetics, and developer-friendly tools for creating new WE methods, including a Python API and resampler module for implementing both binned and “binless” WE strategies.

**Table of Contents/Abstract Image:** For the manuscript “WESTPA 2.0: High-performance upgrades for weighted ensemble simulations and analysis of longer-timescale applications” by Russo *et al*.

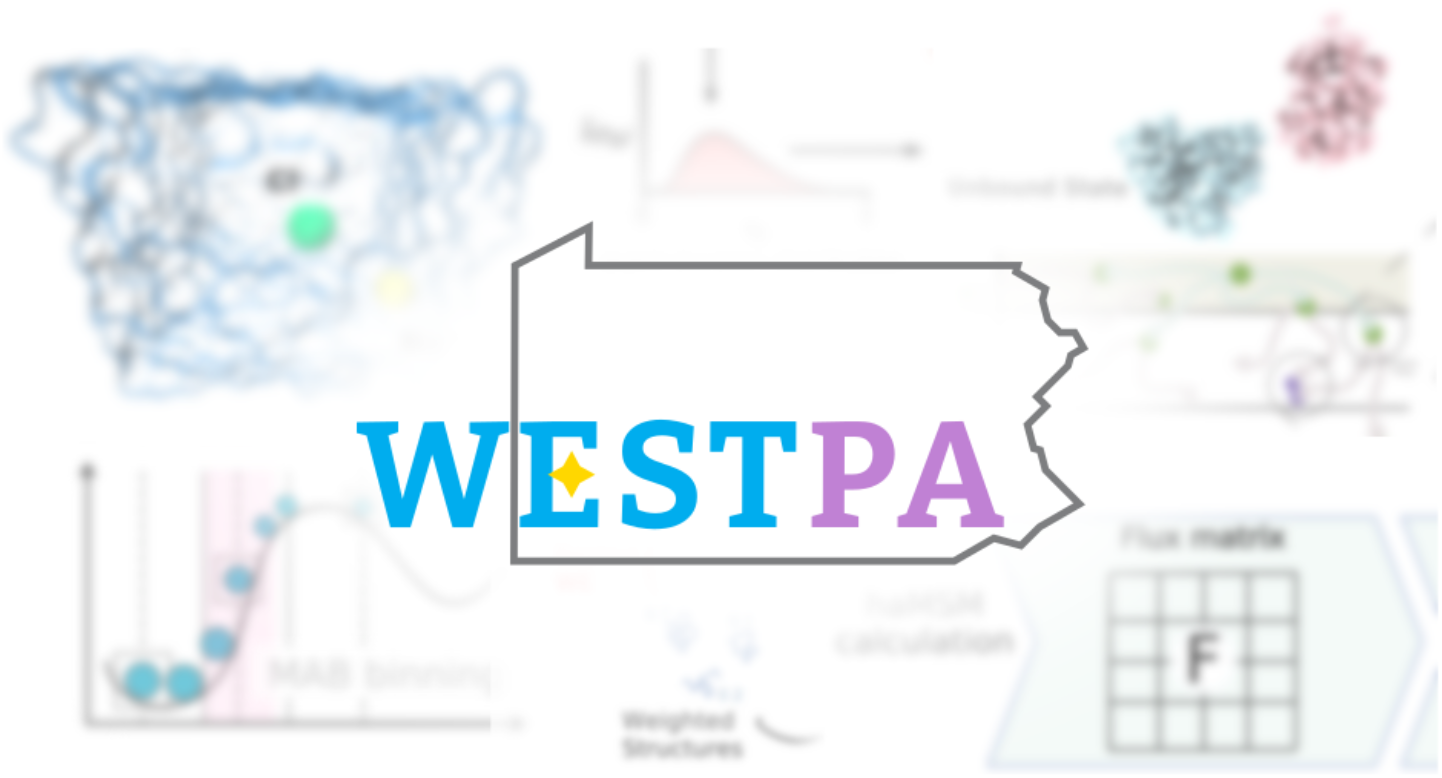

## 1. Introduction

The field of molecular dynamics (MD) simulations of biomolecules arguably is following a trajectory that is typical of mathematical modeling efforts: as scientific knowledge grows, models grow ever more complex and ambitious, rendering them challenging for computation. While early MD simulations focused on single-domain small proteins,^1^ modern simulations have attacked ever larger complexes^2,3^ and even entire virus particles.^4–7^ This trend belies the fact that record-setting small-protein simulations in terms of total simulation time remain limited to the ms scale on special-purpose resources^8^ and to < 100 µs on typical university clusters. These limitations have motivated the development of numerous approaches to accelerate sampling, among which are rigorous path-sampling approaches capable of providing unbiased kinetic and mechanistic observables.^9–18^

Our focus is the weighted ensemble (WE) path sampling approach,^17,19^ which has helped to transform what is feasible for molecular simulations in the generation of pathways for long-timescale processes (> µs) with rigorous kinetics. Among these simulations are notable applications, including atomically detailed simulations of protein folding,^20^ coupled protein folding and binding,^21^ protein-protein binding,^22^ protein-ligand unbinding,^23^ and the large-scale opening of the SARS-CoV-2 spike protein.^24^ The latter is a significant milestone—both in system size (half a million atoms) and timescale (seconds).^24^ Instrumental to the success of the above applications have been advances in not only WE methods, but also software.^24^

Here, we present the next generation (version 2.0) of the most cited, open-source WE software called WESTPA (Weighted Ensemble Simulation Toolkit with Parallelization and Analysis).^25^ WESTPA 2.0 is designed to further enhance the efficiency of WE simulations with high-performance algorithms for: (i) further enhanced sampling via restarting from reweighted trajectories, adaptive binning, and/or binless strategies, (ii) more efficient handling of large simulation datasets, and (iii) analysis tools for estimation of first-passage-time distributions and for more efficient estimation of rate constants. Like its predecessor, WESTPA 2.0 is a highly scalable, portable, and interoperable Python package that embodies the full range of WE’s capabilities, including rigorous theory for any type of stochastic dynamics (e.g., molecular dynamics and Monte Carlo simulations) that is agnostic to the model resolution.^26^ In comparison to other open-source WE packages such as AWE-WQ^27^ and wepy,^28^ WESTPA is unique in its (i) high scalability with nearly perfect scaling out to thousands of CPU cores^24^ and GPUs, and (ii) demonstrated ability to interface with a variety of dynamics engines and model resolutions, including atomistic,^22^ coarse-grained,^29^ whole-cell,^30^ and non-spatial systems models.^31,32^

After a brief overview of the WE strategy (**Section 2**), we describe the organization of WESTPA 2.0 (**Section 3**) and new analysis tools that further expand the capabilities of the software package (**Section 4**). Together, these features greatly facilitate the execution and analysis of WE simulations of even larger systems and/or slower timescales.

## 2. Overview of the weighted ensemble path sampling strategy

The weighted ensemble (WE) strategy enhances the sampling of rare events (e.g., protein folding, binding, chemical reactions) by orchestrating the periodic resampling of multiple, parallel trajectories at fixed time intervals *τ* (**Figure 1**).^17^ The statistically rigorous resampling scheme maintains even coverage of configurational space by replicating (“splitting”) trajectories that have made transitions to newly visited regions and potentially terminating (“merging”) trajectories that have over-populated previously visited regions. The configurational space is typically defined by a progress coordinate that is divided into bins where even coverage of this space is defined as a constant number of trajectories occupying each bin; alternatively, trajectories may be grouped by a desired feature for “binless” resampling schemes.^33^ Importantly, trajectories are assigned statistical weights that are rigorously tracked during resampling; when trajectories are replicated in a given bin, the weights are split among child trajectories and when trajectories are terminated in a probabilistic fashion, the weights are merged with a continued trajectory of that bin. This rigorous tracking ensures that *no bias is introduced into the ensemble dynamics*, enabling direct estimates of rate constants.^26^

**Figure 1.**
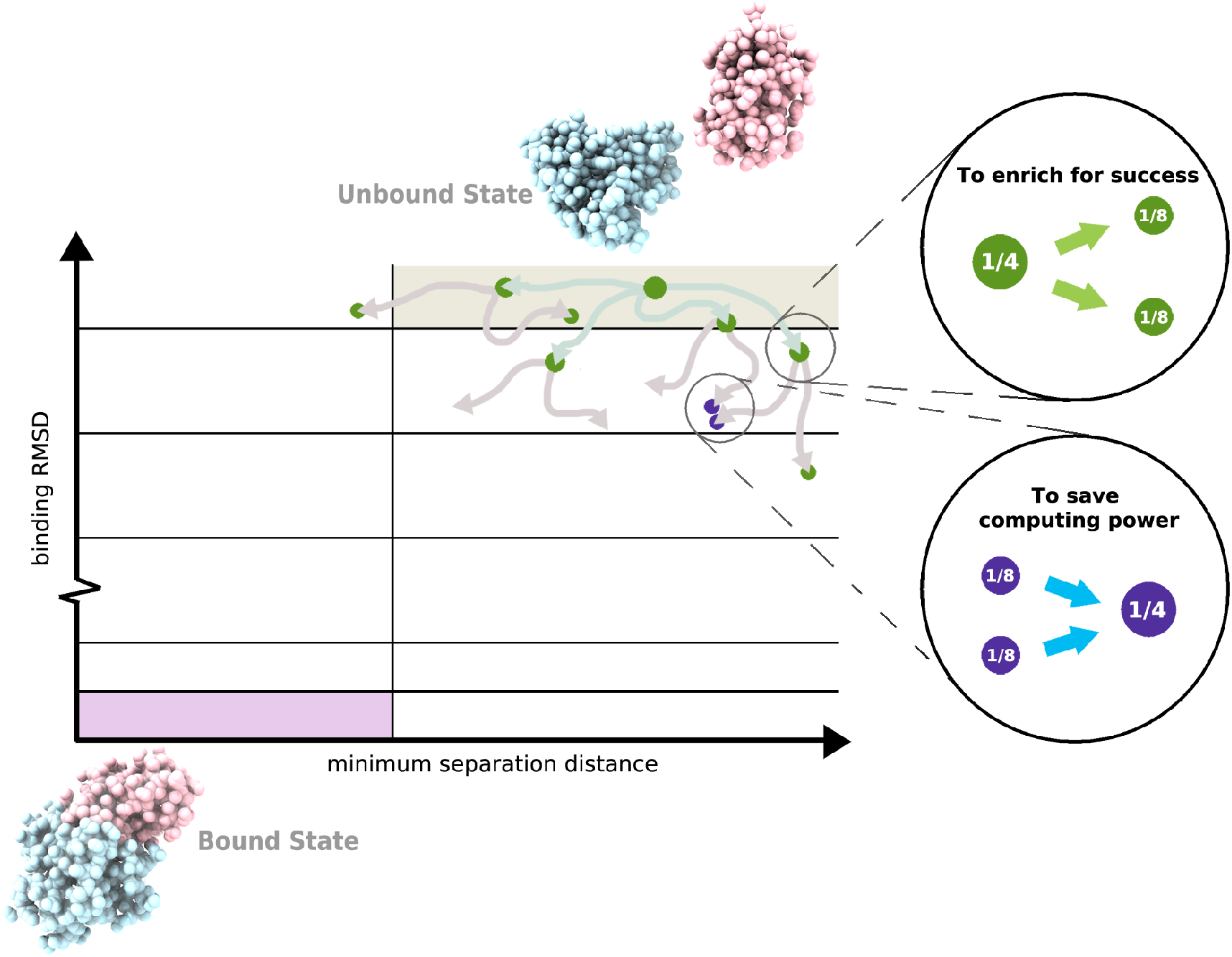
Basic weighted ensemble protocol. As illustrated for the simulation of a protein-protein binding process, a two-dimensional progress coordinate is divided into bins with the goal of occupying each bin with a target number of four trajectories. Four equally weighted trajectories are initiated from the unbound state and subjected to a resampling procedure at periodic time intervals τ: (i) to enrich for success, trajectories that make transitions to less-visited bins are replicated to generate a target of four trajectories in those bins, splitting the weights evenly among the child trajectories (green spheres), and (ii) to save computing time, the lowest-weight trajectories in bins that have exceeded four trajectories are terminated, merging their weights with those of higher-weight trajectories in those bins (purple spheres). Spheres are sized according to their statistical weights.

WE simulations can be run under equilibrium or non-equilibrium steady state conditions. To maintain non-equilibrium steady state conditions, trajectories that reach the target state are “recycled” back to the initial state, retaining the same statistical weight.^34^ The advantage of equilibrium WE simulations over steady-state WE simulations is that the target state need not be strictly defined in advance since no recycling of trajectories at the target state is applied.^35^ On the other hand, steady-state WE simulations have been more efficient in yielding successful pathways and estimates of rate constants. Equilibrium observables can be estimated from either equilibrium WE simulations or the combination of two non-equilibrium steady-state WE simulations in opposite directions when history information is taken into account.^35^

## 3. Organization of WESTPA 2.0

Below, we present the organization of WESTPA 2.0, beginning with code reorganization to facilitate software development (**Section 3.1**) and then proceeding to a description of a Python API for setting up, running, and analyzing WE simulations (**Section 3.2**); a minimal adaptive binning mapper (**Section 3.3**); a generalized resampler module that enables the implementation of both binned and binless schemes (**Section 3.4**); and an HDF5 framework for more efficient handling of large simulation datasets (**Section 3.5**).

### 3.1. Code reorganization to facilitate software development

The WESTPA 2.0 software is designed to facilitate the maintenance and further development of the software according to established and emerging best practices for Python development and packaging. The code has been consolidated and reorganized to better indicate the role of each module (**Figure 2**). The software can now be installed as a standard Python package using pip or by running setup.py. The package will continue to be available through Conda via conda-forge, which streamlines the installation process by enabling WESTPA and all software dependencies to be installed at the same time. We have implemented automated GitHub Actions for continuous integration testing and code quality checks using the Black Python code formatter as a pre-commit hook, alongside flake8 for non-style linting. Templates are provided for GitHub issues and pull requests. Both user’s and developer’s guides are available on the GitHub wiki along with Sphinx documentation of key functions with autogenerated docstrings. Further support will continue to be provided through WESTPA users’ and developers’ email lists hosted on Google Groups (linked on https://westpa.github.io).

**Figure 2.**
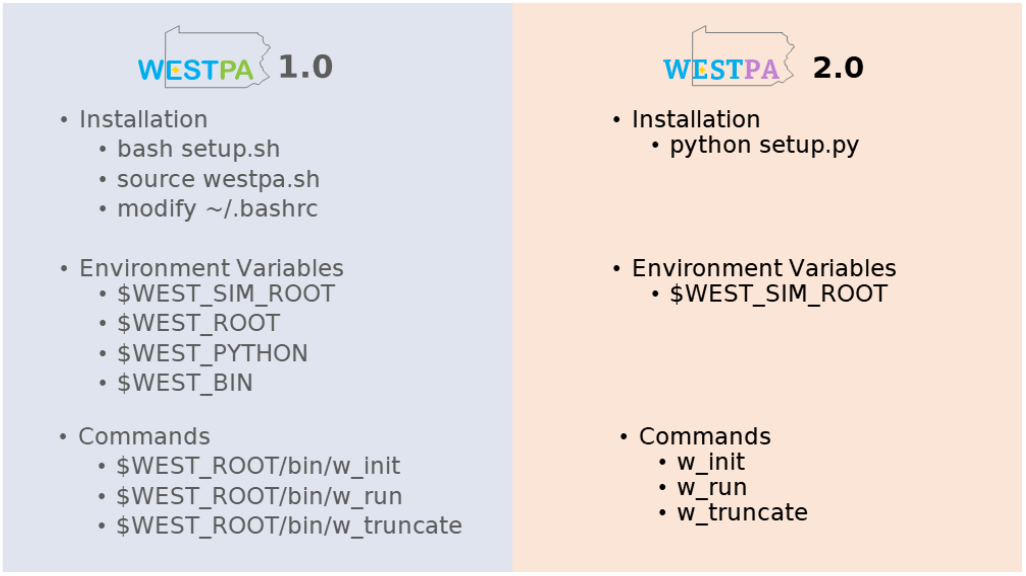
Reorganization of WESTPA 1.0 to WESTPA 2.0. In version 2.0, WESTPA is installed using Python and relies on only a single environment variable such that commands can be called directly through Python. To reflect these changes, we have updated our original suite of WESTPA tutorials for version 2.0 (https://github.com/westpa/westpa_tutorials/tree/westpa-2.0-restruct).^36,37^

### 3.2 Python API for setting up, running, and analysis of WE simulations

To simplify the process of setting up and running WE simulations, WESTPA 2.0 features a Python API that enables the user to execute the relevant commands within a single Python script instead of invoking a series of command-line tools, as previously done in WESTPA 1.0 (**Figure 3A**). This also provides tools for third-party developers to build and develop WESTPA-based applications and plugins, for example, the integration of WESTPA into the cloud-based computing platform, OpenEye Scientific’s Orion;^38,39^ or the haMSM restarting plugin (**Section 4.2**), which uses the results of a WESTPA simulation to perform analysis then restart the simulation based on the results of that analysis.

**Figure 3.**
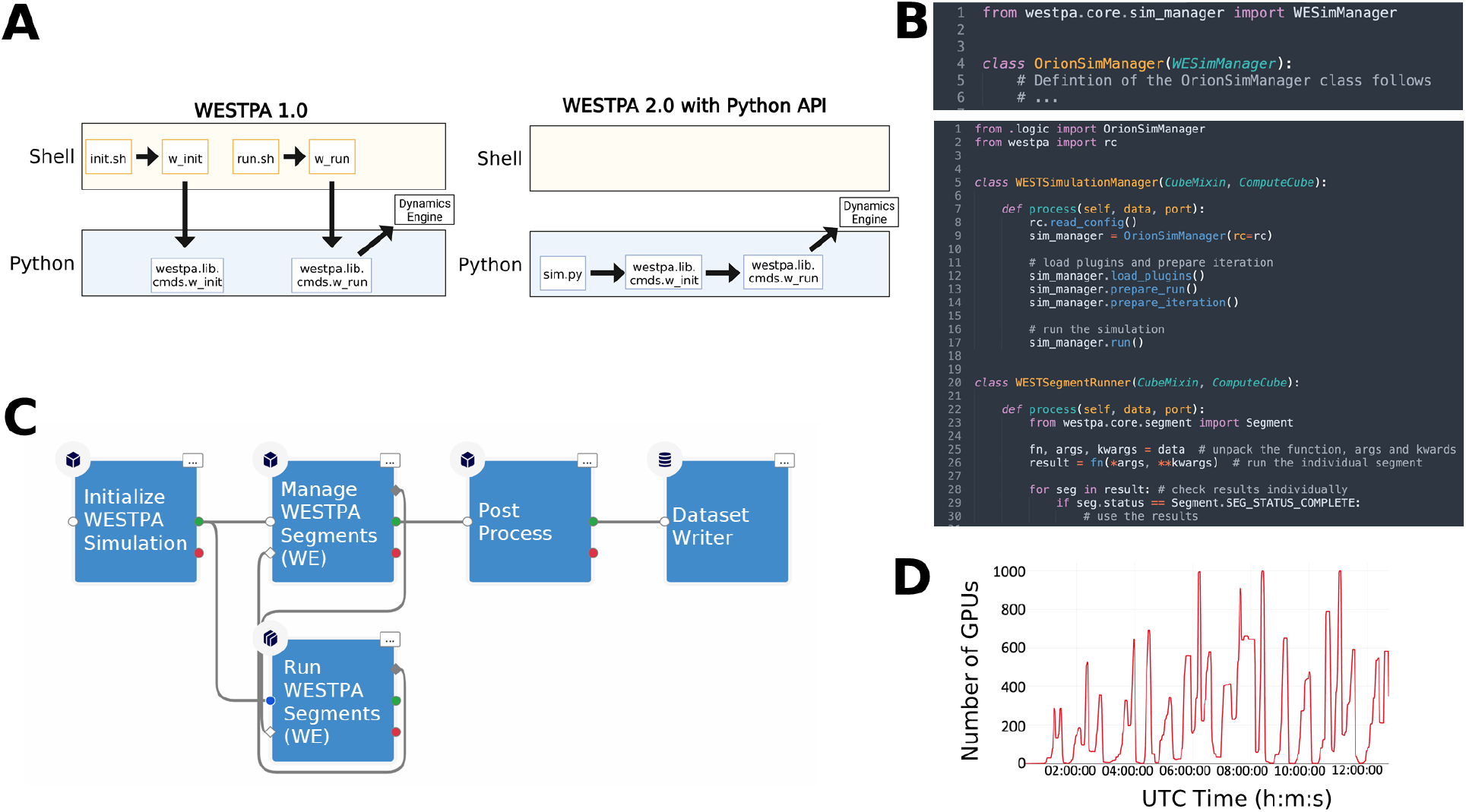
Comparison of workflows for setting up and running WE simulations using WESTPA 1.0 and 2.0, a demonstration of using the Python API for WESTPA 2.0, and GPU performance of the updated API within a cloud computing environment. (A) The Python API in WESTPA 2.0 enables a user to fully define, initialize, and run a WESTPA simulation from within a single Python script (right panel), without needing to invoke command-line utilities required in WESTPA 1.0 (left panel). For backwards compatibility, all original functionality provided in version 1.0 for invoking WESTPA (e.g., `w_init` and `w_run` tools) via shell scripts remains available in WESTPA 2.0. (B) Example of defining a custom simulation manager with the WESTPA 2.0 API (top panel), and using the newly defined simulation manager and WESTPA 2.0 API to programmatically control and run a WE simulation (bottom panel). Here, the WESTSimulationManager class sends work to the WESTSegmentRunner class that unpacks and runs the scripts specified from the WESTPA config file (west.cfg). (C) Example workflow diagram from the Orion user interface using the Python classes constructed from the internal WESTPA APIs presented in **Figure 3B**. Here, a kernel (Initialize WESTPA Simulation) initializes both the `WESTSimulationManager` (Manage WESTPA Segments) and the `WESTSimulationRunner` (Run WESTPA Segments) kernels from **Figure 3B**, which are connected in a cycle to manage splitting and merging. Finally, all data is exported through a Post Process and Dataset Writer kernel for final data processing and storage. (D) Performance of the WESTPA 2.0 API using the `WESTSimulationRunner` class from **Figure 3B** within an Amazon Web Services environment using a combination of numerous `g4dn` instances as a function of wallclock time in Universal Coordinated Time (UTC) units. Here, the per-iteration scaling reaches thousands of GPUs in just under a few hours for a test system of butanol crossing a neat POPC membrane bilayer using the WESTPA 2.0 API with the OpenMM 7.5 MD engine.^41^

**Figure 3B** provides an example of how to programmatically call the WESTPA 2.0 API from the Orion cloud platform, which could in principle be any Python script within any supercomputing or personal computing environment. First, a developer can write any custom simulation or work manager of their choice by subclassing or completely rewriting core WESTPA components (top panel). Second, a workflow can be constructed by invoking a simple set of WESTPA 2.0 Python commands to perform any WE simulation (bottom panel). Typically, a user of the WESTPA 2.0 Python API only needs a handful of API endpoints to perform a complicated simulation protocol. As an example of the power of the simplicity of the Python API, we demonstrate how a workflow can be constructed from defined workflow kernels (**Figure 3C**), and show GPU performance over wall-clock time (in Coordinated Universal Time; UTC) from a drug-like molecule in a membrane permeability simulation (**Figure 3D**). Using the internal API, a user’s simulation can request large amounts of compute resources per iteration. In this case, thousands of GPUs are requested per WE iteration for a simulation of butanol crossing a natural membrane mimetic system (https://github.com/westpa/westpa2_tutorials).^40^

To facilitate the development of custom analysis workflows in cases where more flexibility is required than the existing `w_ipa` analysis tool,^36^ WESTPA 2.0 includes the new `westpa.analysis` Python API. This API provides a high-level view of the data contained in the main WESTPA HDF5 file (“west.h5”), including the trajectory data, reducing the overhead of writing custom analysis code in Python and doing quick, interactive analysis of trajectories (or walkers). The `westpa.analysis` API is built on three core data types: `Run`, `Iteration`, and `Walker`. A `Run` is a sequence of `Iterations`; an `Iteration` is a collection of `Walkers`. Key instance data can be accessed via attributes and methods. For example, a `Walker` has attributes such as the statistical weight (`weight`), progress coordinate value (`pcoords`), starting conformation (`parent`), and child trajectories after replication (`children`), and a method `trace()` to trace its history (as a pure Python alternative to the `w_trace` tool). The API also provides facilities for retrieving and concatenating trajectory segments. These include support for (i) type-aware concatenation of trajectory segments represented by NumPy arrays or MDTraj trajectories, (ii) use of multiple threads to potentially increase performance when segment retrieval is an I/O bound operation, and (iii) display of progress bars. Finally, the API provides a convenience function, `time_average()`, for computing the time average of an observable over a sequence of `Iterations` (e.g., all or part of a `Run`).

### 3.3. A minimal adaptive binning mapper

To automate the placement of bins along a chosen progress coordinate during WE simulation, we have implemented the Minimal Adaptive Binning (MAB) scheme^42^ as an option in the `westpa.core.binning` module. The MAB scheme positions a specified number of bins along a progress coordinate after each resampling interval τ by (1) tagging the positions of the trailing and leading trajectories along the progress coordinate and evenly placing a specified number of bins between these positions, and (2) tagging “bottleneck” trajectories positioned on the steepest probability gradients and assigning these trajectories to their own bins (**Figures 4A-B**). Despite its simplicity, the MAB scheme requires less computing time than manual, fixed binning schemes in surmounting large free energy barriers resulting in more efficient conformational sampling and estimation of rate constants.^42^ To apply the MAB scheme, users specify the `MABBinMapper` option along with accompanying parameters such as the number of bins in the west.cfg file (**Figure 4C**).

**Figure 4.**
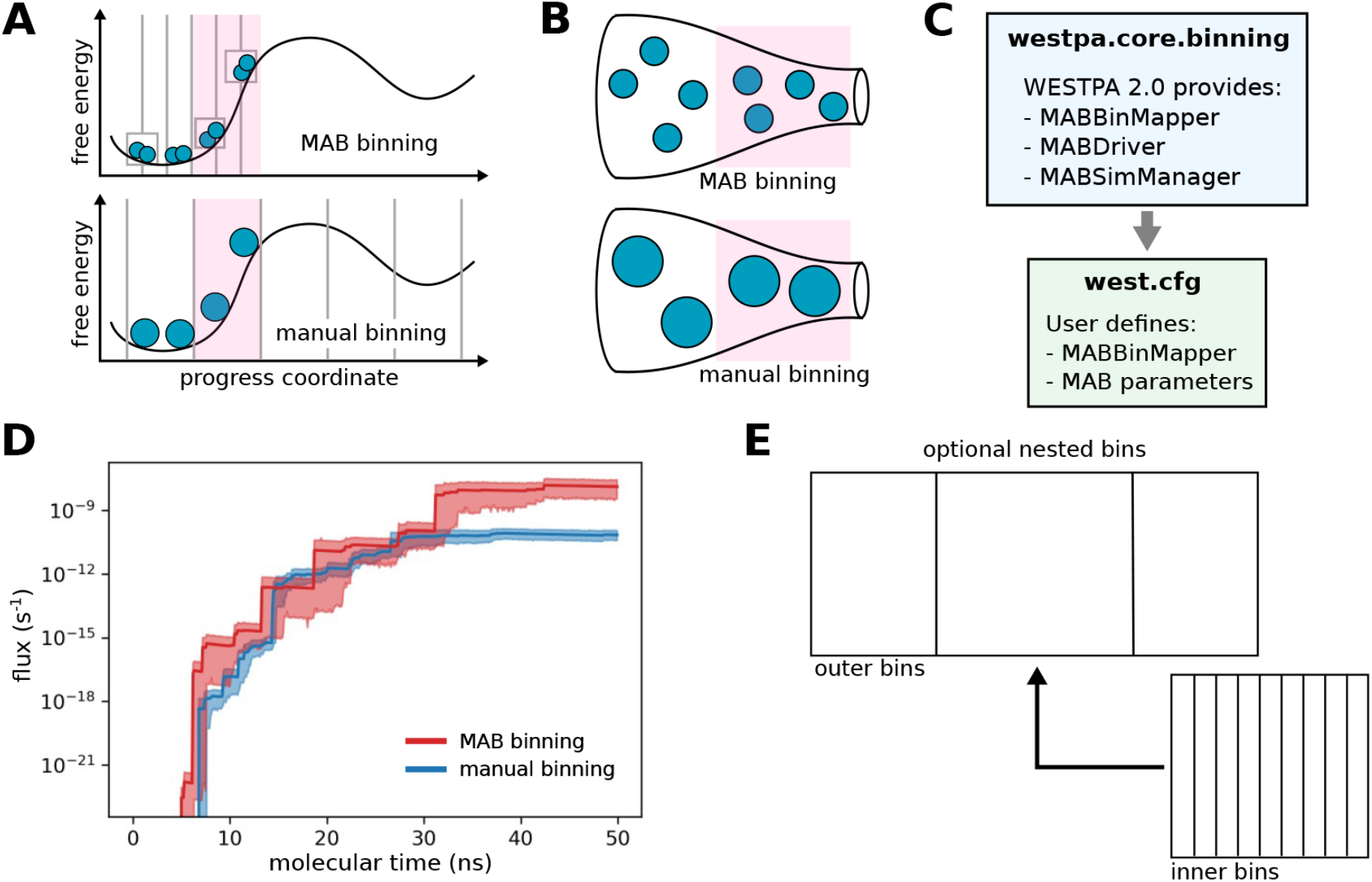
The minimal adaptive binning (MAB) scheme is more efficient in surmounting free energy barriers than manual, fixed binning schemes. (A) Bin positions and trajectories after replication using the MAB scheme vs. a manual binning scheme with the same positions of trajectories (blue circles, sized according to statistical weights) along a chosen progress coordinate and a target of two trajectories per bin. The MAB scheme adaptively positions bins along the progress coordinate by placing equally spaced bins (in this case, three bins, as indicated by solid vertical lines) between the positions of the trailing and leading trajectories along with separate bins (boxes) for these trajectories and a third trajectory in a bottleneck region (pink) along the free energy barrier. (B) Enlarged “bottle” diagrams highlighting the bottleneck region (pink) along with relative positions and weights of trajectories for the MAB and manual binning schemes in panel A). In contrast to the manual binning scheme where trajectories may stall in a bottleneck region, the MAB scheme automatically detects trajectories in this region, replicating these trajectories to enrich for success in surmounting the barrier. (C) MAB-scheme options in the `westpa.core.binning` module and corresponding user-defined options in the west.cfg file. (D) Flux of a drug-like molecule (tacrine) permeating through a neat POPC membrane as a function of molecular time using fixed binning (blue) or adaptive binning (MAB scheme) (red). Solid lines represent mean fluxes and the shaded regions represent 95% confidence intervals. The molecular time is defined as Nτ, where N is the number of WE iterations and τ is the fixed time interval (100 ps) of each WE iteration. Simulations were run using WESTPA 2.0 and the OpenMM 7.5 MD engine.^41^ (E) Schematic of a simple recursive binning case in which closely spaced inner bins are “nested” within a wider outer bin.

**Figure 4D** illustrates the effectiveness of the MAB scheme in enhancing the efficiency of simulating the membrane permeability of a drug-like molecule (tacrine). Relative to a fixed binning scheme, the MAB scheme results in earlier flux of tacrine through a model cellular membrane bilayer (∼5 ns vs. ∼7 ns) and this flux increases more quickly, achieving values that are two orders of magnitude higher for the duration of the test.

The MAB scheme provides a general framework for user creation of more complex adaptive binning schemes.^42^ Users can now specify nested binning schemes in the west.cfg file (**Figure 4E**). To run WESTPA simulations under non-equilibrium steady-state conditions (i.e. with “recycling” of trajectories that reach the target state) with the MAB scheme, users can nest a `MABBinMapper` inside of a `RecursiveBinMapper` bin and specify a target state as the outer bins. Multiple individual `MABBinMappers` can be created and placed at different locations of the outer bins using a recursive scheme, offering further flexibility in the creation of advanced binning schemes.

### 3.4. Generalized resampler module that enables binless schemes

In the original (default) weighted-ensemble resampling scheme, trajectories are split and merged based on a predefined set of bins.^17^ In WESTPA 2.0, we introduce a generalized resampler module that enables users to implement both binned and “binless” resampling schemes, providing the flexibility to resample trajectories based on a property of interest by defining a grouping function. While grouping on the state last visited (e.g., initial or target state) was previously possible using the binning machinery in WESTPA 1.0,^43^ our new resampler module provides a more general framework for creating binless schemes by defining a group/reward function of interest such as nonlinear progress coordinates that may be identified by machine learning techniques. Following others,^45^ the resampler module includes options for (i) specifying a minimum threshold for trajectory weights to avoid running trajectories with inconsequentially low weights, and (ii) specifying a maximum threshold for trajectory weights to avoid a single large-weight trajectory from dominating the sampling, increasing the number of uncorrelated successful events that reach the target state.

As illustrated in **Figure 5**, the implementation of a binless scheme requires two modifications to the default WESTPA simulation: (i) a user-provided group module containing the methods needed to process the resampling property of interest for each trajectory walker, and (ii) updates to the west.cfg file specifying the resampling method in the `group_function` keyword and the attribute in the `group_arguments` keyword.

**Figure 5.**
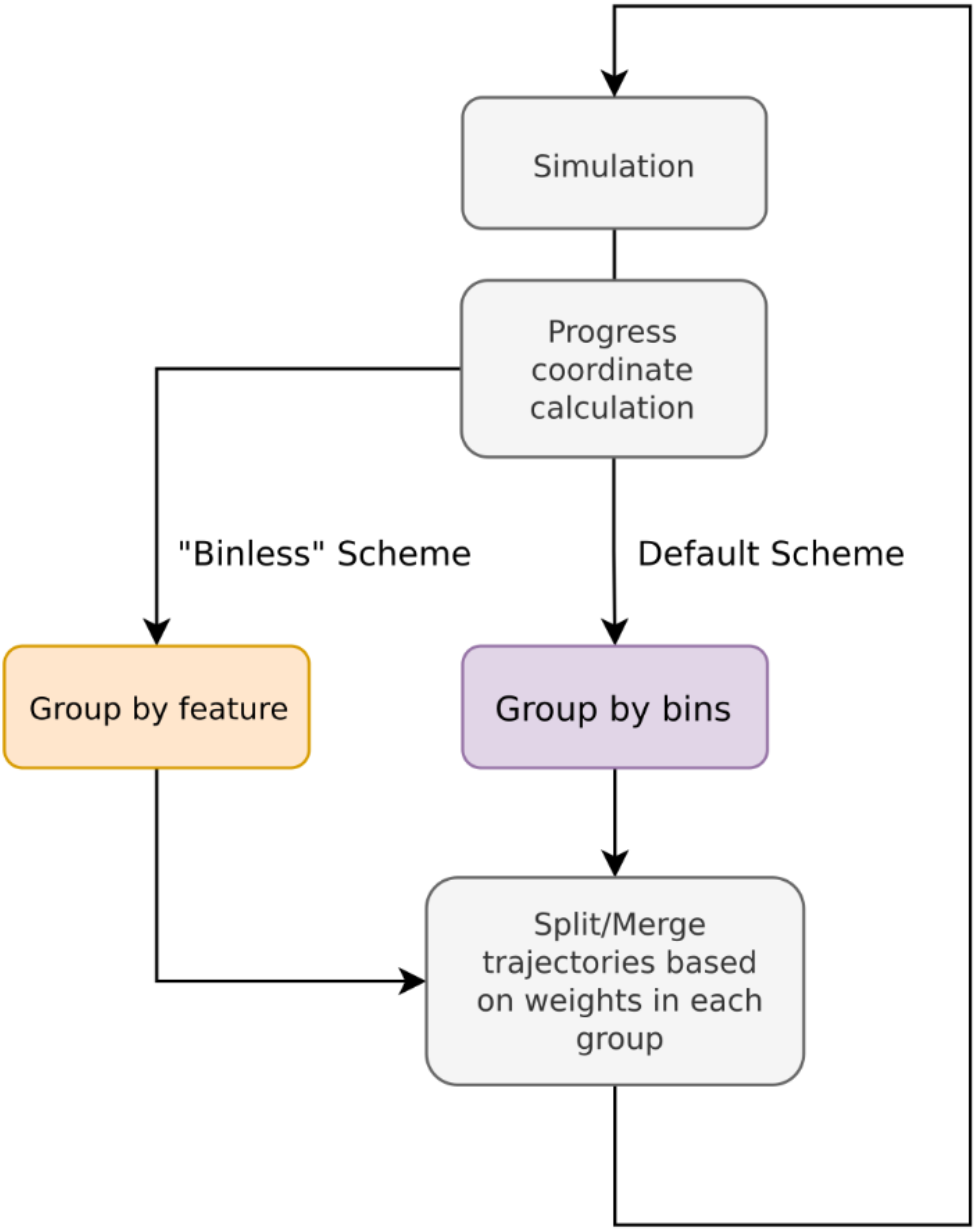
Flowchart for implementing binless resampling schemes in WESTPA 2.0. The implementation involves grouping trajectories by feature (using the `group_function` defined in the `group` module) before splitting and merging. The functionality for positioning bins along a chosen progress coordinate remains available in WESTPA 2.0.

We provide two examples of implementing binless schemes in the westpa-2.0-restruct branch of the WESTPA_Tutorials GitHub repository (https://github.com/westpa/westpa_tutorials/tree/westpa-2.0-restruct).^37^ The basic_nacl_group_by_history example illustrates grouping of trajectory based on its “history”, i.e. a shared parent *N* WE iterations back. The parameter *N* is specified in the keyword hist_length under the `group_arguments` keyword in the west.cfg file. This WESTPA configuration file also specifies the name of the grouping function method, `group.walkers_by_history`, in the `group_function` keyword. In the basic_nacl_group_by_color example, trajectory walkers are tagged based on “color” according to the state last visited. Only walkers that have the same color are merged, thereby increasing the sampling of pathways in both directions. State definitions are declared within the `group_arguments` keyword in the west.cfg file.

### 3.5. HDF5 framework for more efficient handling of large simulation datasets

One major challenge of running WE simulations has been the management of the resulting large datasets, which can amount to tens of terabytes over millions of trajectory files. To address this challenge, we have developed a framework for storing trajectory data in a highly compressed and portable HDF5 file format. The format is derived from the HDFReporter class implemented in the MDTraj analysis suite,^46^ and maintains compatibility with NGLView,^47^ an iPython/Jupyter widget for interactive viewing of molecular structures and trajectories. A single HDF5 file is generated per WE iteration, which includes a link to each trajectory file stored in the main WESTPA data file (west.h5). Thus, the new HDF5 framework in WESTPA 2.0 enables users to restart a WE simulation from a single HDF5 file rather than millions of trajectory files and simplifies data sharing as well as analysis. The dramatic reduction in the number of trajectory files also eliminates potentially large overhead from the filesystem that results from the storage of numerous small files. For example, a 53% overhead has been observed for a 7.5-GB dataset of 103,260 trajectory files generated from NTL9 protein folding simulations (**Figure 9**), occupying 11.5 GB of actual disk storage on a Lustre filesystem.

To test the effectiveness of the HDF5 framework in reducing the amount of data storage required for WE simulations, we applied the framework to a set of three independent WE simulations of Na^+^/Cl^-^ association and one WE simulation involving p53 peptide conformational sampling (**Figures 6A-B**). Our results revealed 27% and 85% average reduction in the total size of trajectory files generated during the Na^+^/Cl^-^ association and p53 peptide simulations, respectively, relative to WESTPA 1.0. Given a fixed number of bins, the sizes of per-iteration HDF5 files were also shown to converge as the simulation progresses (**Figures 6C-D**), suggesting that the storage of trajectory data by iteration not only facilitates the management of the data but also yields files of roughly equal sizes. The difference in the reduction efficiency that we observed between the Na^+^/Cl^-^ and p53 peptide systems can be attributed to differences in the simulation configurations including the format of the output trajectories, restart files and other factors such as the verbosity of logging.

**Figure 6.**
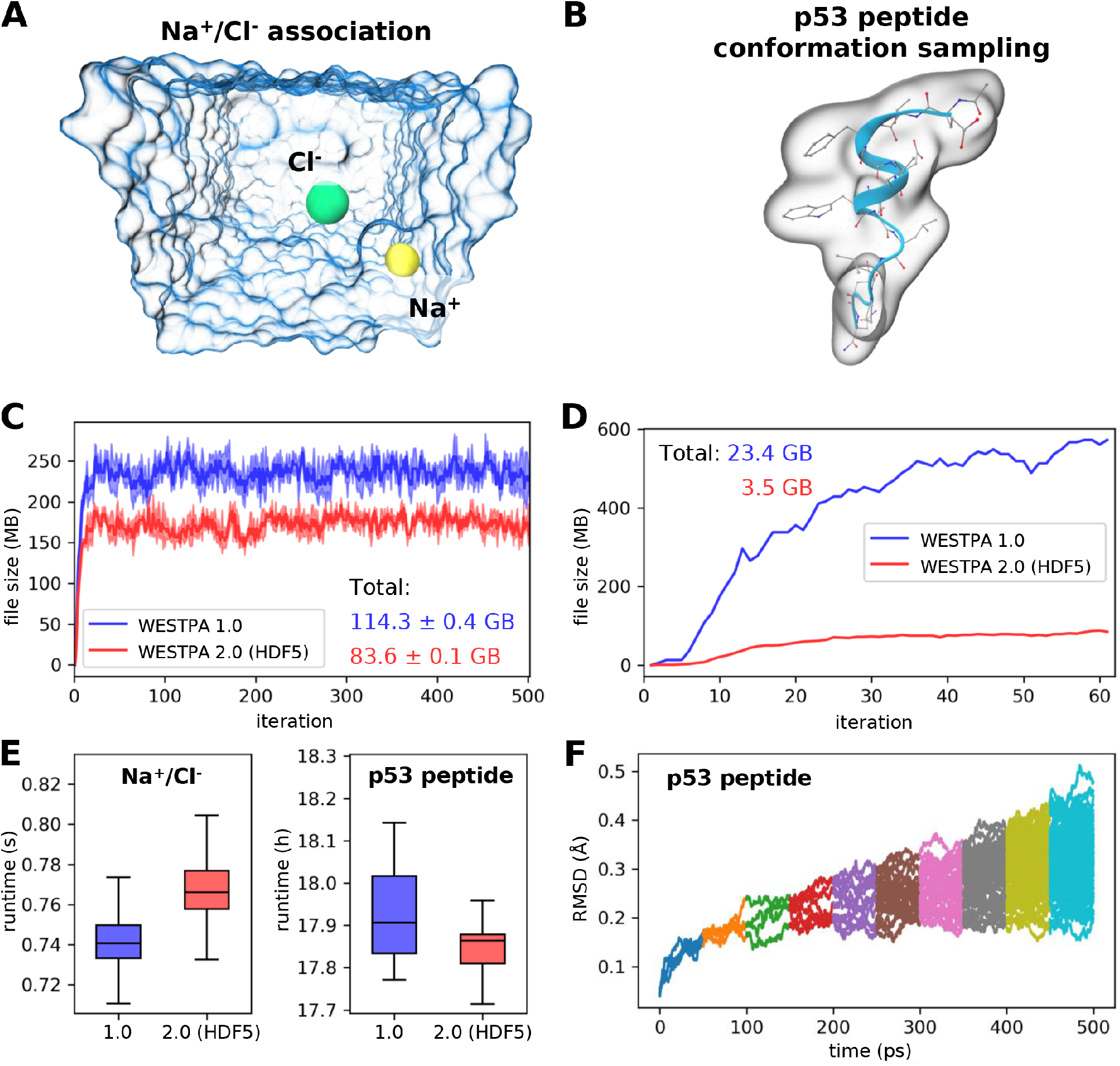
Demonstration of the usage of the HDF5 framework for two example systems. (A) Na+/Cl- association simulation where Na+ (*yellow sphere*) and Cl- (*green sphere*) ions were solvated in explicit water (*blue transparent surface*). The distance between the two ions serves as the progress coordinate. (B) Conformational sampling of a p53 peptide (residues 17-29) in generalized Born implicit solvent using a progress coordinate consisting of the heavy-atom RMSD of the peptide from its MDM2-bound conformation.^21^ The molecular surface of the p53 peptide is rendered as a transparent surface, with both the secondary (*blue ribbon*) and atomic structures overlaid. (C) Comparison of file sizes of per-iteration HDF5 files for the Na^+^/Cl^-^ association simulation as a function of the WE iteration using WESTPA 1.0 and 2.0 with the HDF5 framework. The result was obtained from three independent simulations where the *solid curves* show the mean file sizes, while the *light bands* show the standard deviations. (D) Same comparison as panel C for a single simulation of the p53 peptide, hence no error bars are shown. (E) Comparison of wall-clock runtimes normalized by the number of trajectory segments per WE iteration using WESTPA 1.0 and 2.0 with the HDF5 framework option turned on. (F) Time-evolution of the heavy-atom RMSD of the p53 peptide from its MDM2-bound conformation by trajectories obtained using WESTPA’s analysis tools. Colors represent RMSDs obtained from different iterations. WESTPA simulations of Na^+^/Cl^-^ association and the p53 peptide were run using the OpenMM 7.5 MD engine.^41^

Our tests revealed that the additional steps introduced by the HDF5 framework for managing trajectory coordinate and restart files did not have any significant impact on the WESTPA runtime (**Figure 6E**), which is normalized by the number of trajectory segments per WE iteration given that the evolution of bin occupancies by trajectories can vary among different runs due to the stochastic nature of the MD simulations (after 60 iterations, the WESTPA 1.0 run occupied six more bins than the WESTPA 2.0/HDF5 run). This variation in the bin occupancy is unlikely to be affected by the HDF5 framework since it simply manages the trajectory and restart files and does not alter how the system is simulated. The differences in bin occupancies/total number of trajectories may also partially contribute to the large reduction in per-iteration file sizes for the HDF5 run observed in **Figure 6D** of the p53 peptide. However, the majority of this file-size reduction results from efficient HDF5 data compression of trajectory coordinates, restart, and log files. Finally, trajectory data saved in the HDF5 files can be extracted and analyzed easily using MDTraj in combination with our new analysis framework presented in **Section 3.2** (**Figure 6F**).

## 4. Analysis tools

WESTPA 2.0 features new analysis tools for estimating rate constants more efficiently using the distribution of “barrier crossing” times (**Section 4.1**), accelerating convergence using a history-augmented Markov state model to reweight trajectories (**Section 4.2**), and estimating the distribution of first passage times (**Section 4.3**).

### 4.1. The RED scheme for rate-constant estimation

To more efficiently estimate rate constants from WE simulations, we have implemented the Rates from Event Durations (RED) scheme as an analysis tool called `w_red` in the WESTPA 2.0 software. The RED scheme exploits the transient ramp-up portion of a WE simulation by incorporating the probability distribution of event durations (or “barrier crossing” times) from a WE simulation as part of a correction factor (**Figure 7A**).^49^ The correction factor accounts for the systematic error that results from statistical bias toward the observation of events with short durations and reweights the event duration distribution accordingly. When applied to an atomistic WE simulation of a protein-protein binding process, the RED scheme is >25% more efficient than the original WE scheme^17^ in estimating the association rate constant (**Figure 7B**).^49^

**Figure 7.**
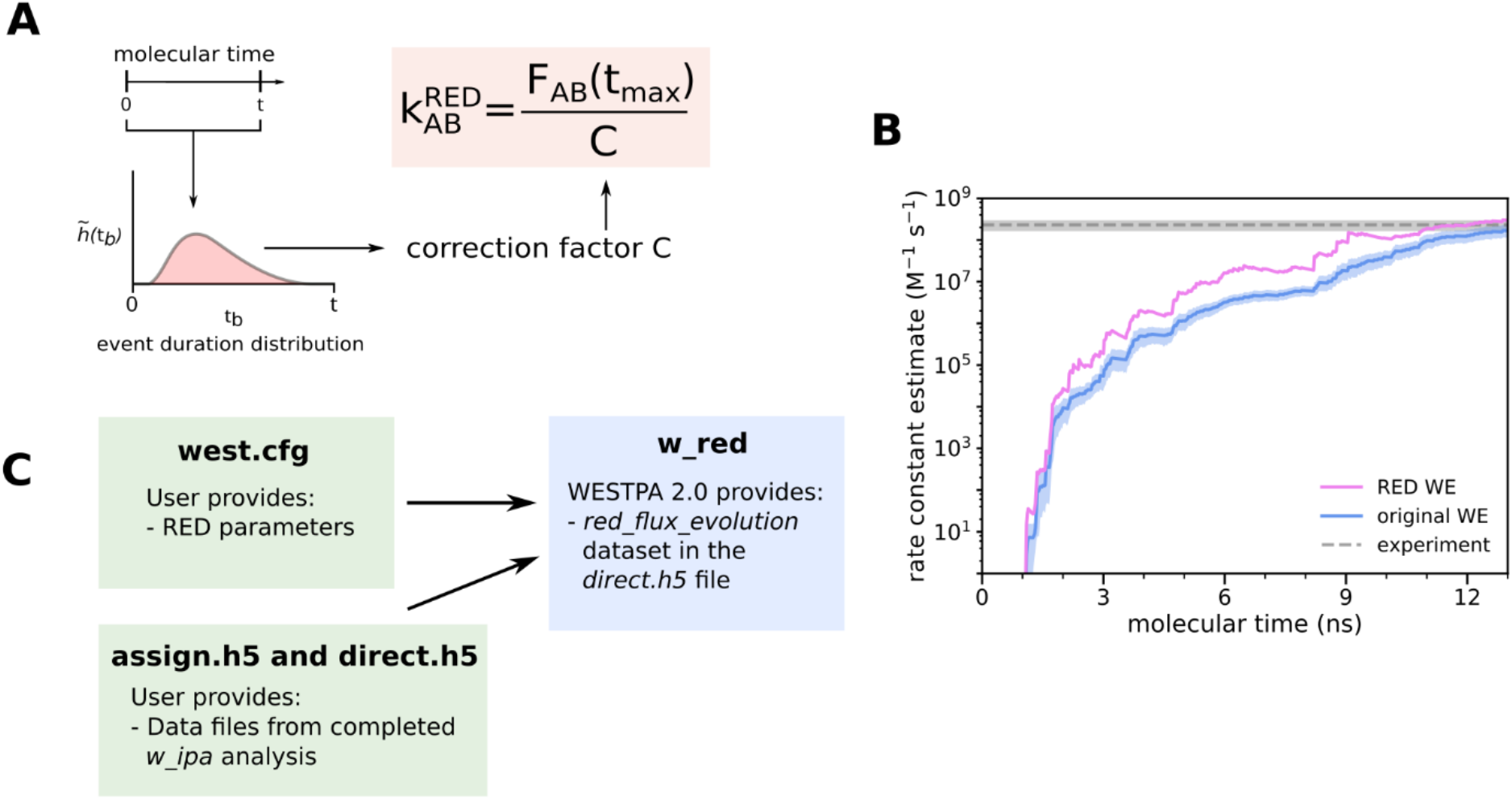
The RED scheme for more efficient rate-constant estimation. (A) Schematic illustrating the RED scheme, which incorporates the distribution of event durations as part of a correction factor for rate-constant estimates that accounts for statistical bias toward the observation of events with short durations. (B) Application of the original and RED schemes to estimate the associate rate constant of a protein-protein binding process involving the barnase and barstar proteins as a function of molecular time in a WE simulation. The molecular time is defined as Nτ, where N is the number of WE iterations and τ is the fixed time interval (20 ps) of each WE iteration. Simulations were previously run using WESTPA 1.0 with the GROMACS 4.6.7 MD engine.^48^ (C) A schematic illustrating how users can generate a dataset for calculating the RED-scheme correction factor from simulation data stored in the analysis HDF5 files and apply the correction factor to the rate-constant estimate using the new `w_red` tool.

The code for estimating rate constants using the RED scheme takes as input the assign.h5 files and direct.h5 files generated by the `w_ipa` analysis tool. Users then specify in the analysis section of the west.cfg file which analysis scheme `w_red` should analyze along with the initial/final states and the number of frames per iteration. Executing `w_red` from the command line, will output the corrected flux estimates as a new dataset called red_flux_evolution to the users’ existing direct.h5 file (**Figure 7C**). The RED rate-constant estimates can then be accessed through the Python h5py module and plotted vs. time to assess the convergence of the estimates. To estimate uncertainties in observables calculated from a small number of trials (i.e. number of independent WE simulations), we recommend using the Bayesian Bootstrap approach.^17,50^ If it is not feasible to run multiple independent simulations with a certain system due to either system size or the timescale of the process of interest, a user can apply a Monte Carlo bootstrapping approach to a single simulation’s RED rate constant estimate.

### 4.2. A history-augmented Markov State Model (haMSM) restarting plugin

History-augmented Markov state models (haMSMs) provide a powerful tool for estimation of stationary distributions and rate constants from transient, unconverged WE data.^51^ Thus, the approach has a similar motivation to the RED scheme.^49^ In haMSM analysis instead of discretizing trajectories via the WE bins used by WESTPA, as in the `WESS`/`WEED` reweighting plugins,^34,35^ a much finer and more numerous set of ‘microbins’ is employed to calculate steady-state properties with higher accuracy. These estimates, in turn, can be used to start new WE simulations from a steady-state estimate, accelerating convergence of the simulation.^50^ The new plugin provides a streamlined implementation of the restarting protocol that runs automatically as part of a WESTPA simulation, a capability which did not previously exist.

The `msm_we` package provides a set of analysis tools for using typical WESTPA HDF5 output files, augmented with atomic coordinates, to construct an haMSM. A nearly typical MSM model-building procedure^53^ is used (**Figure 8**): WE trajectories are discretized into clusters (microbins) and transitions among microbins are analyzed. However, instead of reconstructing entire trajectories, the `msm_we` analysis computes the flux matrix by taking each weighted parent/child segment pair, extracting and discretizing one frame from each, and measuring flux between them - i.e. the weight transferred.

**Figure 8.**
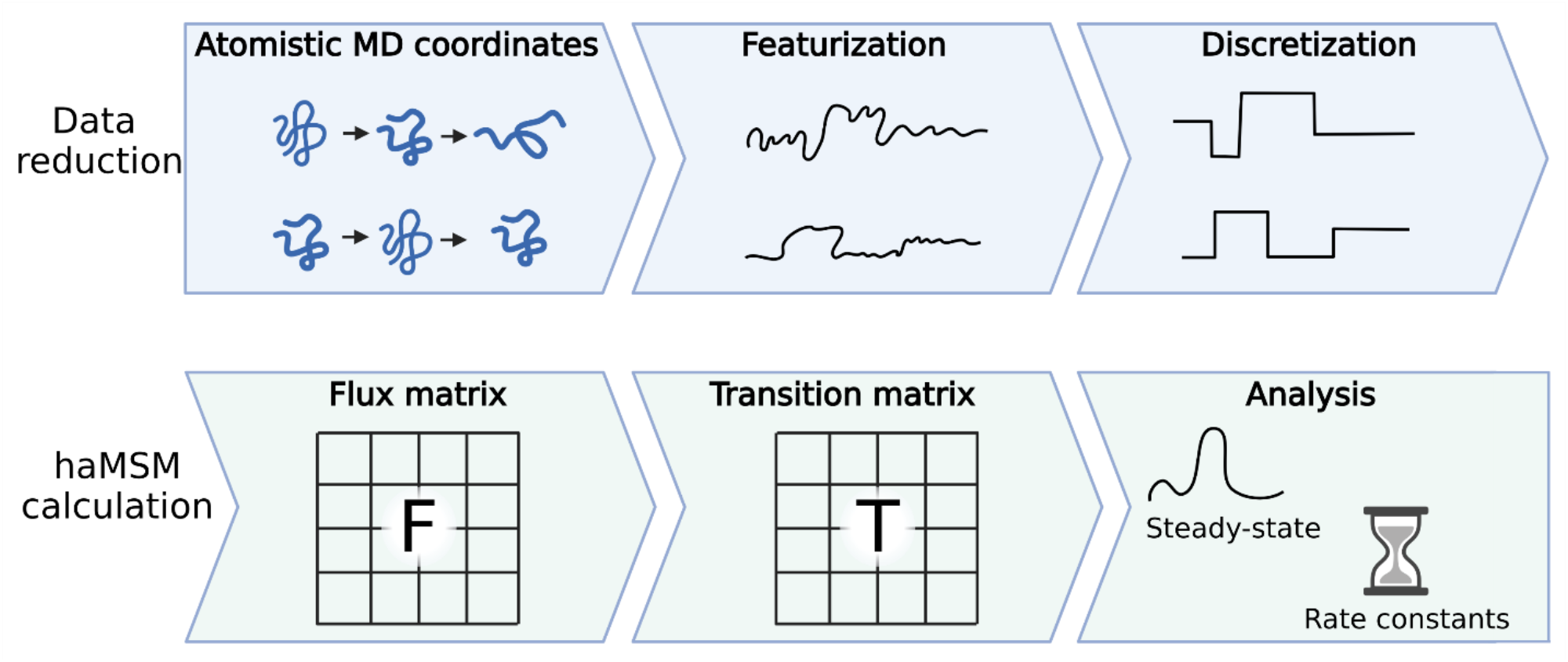
Workflow for constructing an haMSM from trajectories. First, the atomistic trajectories are featurized and discretized. The flux matrix is then computed by computing fluxes between discrete states. The flux matrix is row-normalized into a transition matrix. Estimates of steady-state populations and rate constants are obtained from analysis of the transition matrix.^52^

The haMSM restarting plugin in WESTPA 2.0 makes use of the analysis tools provided by `msm_we` to incorporate this functionality directly into WESTPA. It manages running a number of independent simulations, initialized from some starting configuration, and augments their output HDF5 with the necessary atomic coordinates. Data from all independent runs are gathered and used to build a single haMSM. Stationary probability distributions and rate constants are estimated from this haMSM.

This plugin can be used to start a set of new WE simulation runs, initialized closer to steady-state (**Figure 9**). The haMSM and the WE trajectory data are used to build a library of structures and their associated steady-state weights. These are used to initiate a new set of independent WE runs, which should start closer to steady-state and thus converge more quickly. The process can be repeated iteratively, as shown in **Figure 9A**. The result of this restarting procedure is shown in **Figure 9B**. For challenging systems, the quality of the haMSM will greatly affect the quality of the steady-state estimate. A further report is forthcoming on strategies for building high-quality haMSMs.

**Figure 9.**
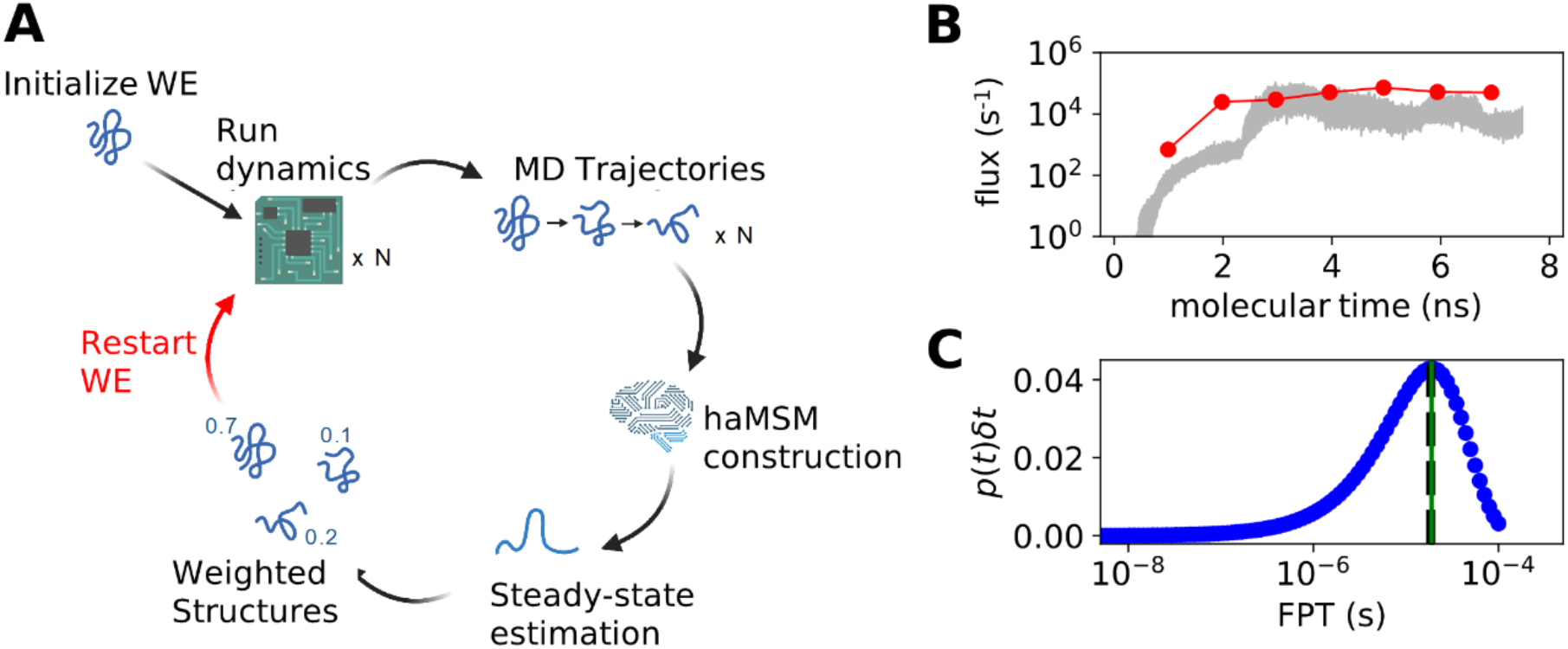
Application of haMSM restarting plugin to the ms folding process of the NTL9 protein. (A) Diagram of the haMSM restarting plugin’s functionality. (B) Example of restarting plugin functionality in accelerated convergence of NTL9 folding rate constants from a WESTPA 2.0 simulation using the AMBER 16 MD engine.^54^ haMSM estimates at restarting points are shown as dots, WE direct fluxes are shown as red lines, and a 95% credibility region from direct WE is shown in gray. (C) Distribution of first passage times for NTL9 folding from the haMSM built at the final restart of the simulation in **Figure 9B**. The weighted average of the blue FPT distribution is shown in black dashed, and the MFPT estimate from the haMSM’s steady-state estimate is shown in green.^52^

To use this plugin, users must specify a function that ingests coordinate data and featurizes the data. Dimensionality reduction may be performed on this featurized data. An effective choice of featurization provides a more granular structural description of the system without including a large number of irrelevant coordinates that add noise without adding useful information. For example, a limited subset of the full atoms such as only alpha-carbons, or even a strided selection of the alpha carbons, may be sufficient to capture the important structural information. Choosing a featurization based on rotation-invariant distances, such as pairwise atomic distances instead of atomic positions, can also help capture structural fluctuations without sensitivity to large-scale motion of the entire system.

To validate convergence of the restarted simulations, a number of independent replicates of the restarting protocol should be performed. These replicates should demonstrate both stability in flux estimates across restarts, and relatively constant-in-time direct fluxes within the restarts. If limited to a single replicate, agreement between the haMSM flux estimate and the direct flux should also be validated.

### 4.3. Estimating first-passage-time distributions

First passage times (FPTs) and their mean values (MFPTs) are key kinetics quantities to characterize many stochastic processes (from a macrostate to another) in chemistry and biophysics such as chemical reactions, ligand binding and unbinding, protein folding, diffusion processes of small molecules within crowded environments. WE simulations, via the Hill relation, provide unbiased estimates of the MFPT directly once steady is reached^34^ or indirectly via non-Markovian haMSM analysis,^35^ but mathematically rigorous estimation of the FPT distribution is not available and has been a challenge for WE simulation. Suárez and coworkers, however, have shown that the FPT distributions estimated from haMSM models provide semi-quantitative agreement with unbiased reference distributions in different systems.^55^ Details on building history-augmented MSMs are described above in **Section 4.2** and more information can be found in the references.^35,55^

Here, we extend and strengthen earlier FPT distribution analysis from WE data. The original code for calculating FPT distribution was published on a separate GitHub repository (https://github.com/ZuckermanLab/NMpathAnalysis).^56^ Recently we reorganized and refactored the code in class hierarchical structures: a base class (`MatrixFPT`) for calculating MFPTs and FPTs distribution using a general transition matrix as an input parameter, and two derived classes (`MarkovFPT` and `NonMarkovFPT`) using transition matrices from Markovian analysis and non-Markovian analysis such as haMSM in **Section 4.2** respectively. The updated code has been merged into the `msm_we` package described in **Section 4.2** along with some updates on building transition matrix from classic MD simulation trajectories.

The new code enables robust estimation of the FPT distribution. **Figure 9C** shows the non-Markovian estimation of the FPT distribution of transitions between macrostate A and B from the WE simulation of NTL9 protein folding.

## 5. Summary

WESTPA is an open-source, high-scalable, interoperable software package for applying the weighted ensemble (WE) strategy, which greatly increases the efficiency of simulating rare events (e.g., protein folding, protein binding) while maintaining rigorous kinetics. The latest WESTPA release (version 2.0) is a substantial upgrade from the original software with high-performance algorithms enabling the simulation of ever more complex systems and processes and implementing new analysis tools. WESTPA 2.0 has also been reorganized into a more standard Python package to facilitate code development of new WE algorithms, including binless strategies. With these features available in the WESTPA toolbox, the WE community is well-poised to take advantage of the latest strategies for tackling major challenges in rare-events sampling, including the identification of slow coordinates using machine learning techniques,^57,58^ and the interfacing of the WE strategy with other software involving complementary rare-event sampling strategies (e.g., OpenPathSampling,^59,60^ SAFFIRE,^61^ and ScMile^62^) and analysis tools (e.g., LOOS,^63,64^ MDAnalysis,^65,66^ and PyEmma^67^). WESTPA has also been interfaced with OpenEye Scientific’s Orion platform^39^ on the Amazon Web Services cloud computing facility. We hope that the above new features of WESTPA will greatly facilitate efforts by the scientific community to tackle grand challenges in the simulation of rare events in a variety of fields, including the molecular sciences and systems biology.

## Notes

The authors declare the following competing financial interest(s). L.T.C. is a current member of the Scientific Advisory Board of OpenEye Scientific and an Open Science Fellow with Roivant Sciences. S.Z., J.P.T., J.X., and D.N.L. are employees of OpenEye Scientific.

## Acknowledgments

This work was supported by an NIH grant (R01 GM115805) to L.T.C. and D.M.Z.; NSF grants (CHE-1807301 and MCB-2112871) to L.T.C.; a MolSSI Software Fellowship to J.D.R.; and a University of Pittsburgh Andrew Mellon Graduate Fellowship to A.T.B. Computational resources were provided by the University of Pittsburgh’s Center for Research Computing, by OpenEye Scientific via compute instances sourced from Amazon Web Services, and by the Advanced Computing Center at Oregon Health and Science University. We thank David Aristoff, Gideon Simpson, Forrest York, Darian Yang, and Alan Grossfield for helpful discussions.

